# Jaguar Conservation Units as Effective Tool in a Payment for Ecosystem Services Scheme for Apex Predator

**DOI:** 10.1101/2025.08.29.673106

**Authors:** Leandro Silveira, Anah Tereza de Almeida Jácomo, Renato Alves Moreira, José Alexandre Felizola Diniz-Filho, Lucas Jardim, Everton B. P. Miranda, Wagner Fischer, Tiago Jácomo, Marcus Vinicius Aguiar Macedo, Erich Raphael Masson, Guilherme Fernandes Ferreira Tavares, Juliano Baiocchi Villa-Verde de Carvalho, Giselle Bastos Alves

## Abstract

This paper proposes a redefinition of Jaguar Conservation Units, emphasizing the role of private rural properties in jaguar populations in Brazil while mitigating conflicts with the species and allowing a more effective implementation of payment for the species’ environmental services. Jaguars prey domestic livestock, leading to retaliatory killings by ranchers. In this context, the study aims to analyze the environmental and economic parameters associated with the ecosystem services provided by jaguars. The ultimate goal is to develop a regional Payment for Environmental Services (PES) scheme to support jaguar conservation by promoting forest code compliance, maintaining habitat connectivity, and compensating cattle predation, using data from Mato Grosso do Sul (MS), Brazil. Our findings demonstrate the advantages of implementing such program and reveal that the cost of maintaining jaguar populations through a PES, with funds from several independent sources, would be feasible and important for one of the largest jaguar populations in Brazil.

**HIGHLIGHTS:**

- Analysis of environmental and economic parameters linked to jaguar ecosystem services.
- Development of a regional Payment for Environmental Services (PES) scheme in Mato Grosso do Sul.
- Evidence of the feasibility and importance of PES to maintain one of the largest jaguar populations in Brazil.

## 1. INTRODUCTION

The jaguar (*Panthera onca* L.) is the largest felid in the Americas, standing as one of the most iconic predators in Earth’s biodiversity. Once seen through mythical lenses, jaguars are now valued ecologically, morally, and economically, though still viewed as threats (Franco & Gonçalves, 2020; Rabinowitz, 2014). As a keystone and flagship species, they are a conservation priority in the Neotropics (UNDP et al., 2018).

Jaguars historically ranged from Argentinean Patagonia to the southern US. Sanderson et al. (2002; see also Nijhawan, 2012), provided the first overview of the conservation status of jaguars at a continental scale, establishing the Jaguar Conservation Units (JCUs), representing diverse habitats and potentially distinct populations. JCUs were defined as areas with suitable habitats and known to encompass large viable populations that would be potentially self-sustainable in the next 100 year, and your relative importance was subsequently evaluated in terms of their size, connectivity, habitat quality, poaching of jaguars and its prey, and population status, by considering the main threats to long-term persistence of these population.

Thus, the reasoning underlying the definition of JCUs is that prioritizing conservation actions within these areas would increase the potential for species’ persistence in the long run, mitigating the rapid loss of its geographic extent due to several threats. Indeed, in Brazil, the current extent of jaguars has been significantly reduced to at least 54% of its historical range, primarily due to the conversion of natural habitats for agricultural and ranching purposes that strongly affect abundance and extent of large carnivores (Jędrzejewski et al., 2023).

If we subtract the governmental protected areas under strict protection from the current jaguar geographic range (JGR), we learn that approximately 95% of its distribution is under human interference. Therefore, note that despite the number of JCUs theoretically defined for Brazil, they are mostly dependent on the private sector to make them effective in jaguar conservation. Thus, these figures open many interesting questions on the definition and meaning of JCUs. While the concept of JCUs is valuable, implementing it in practice is challenging. Creating the extensive reserves necessary to support viable jaguar populations is often hindered by competition for land, the ongoing expansion of agriculture and pasture, and the fact that many of these areas are already occupied by such activities. In this context, Nature-Based Solutions and Payment for Environmental Services (PES) would provide a concrete framework to make JCUs effective units for jaguar conservation (i.e., Albert et al., 2017; Pereira et al., 2023).

Indeed, operationally, various initiatives across the jaguar’s range offer case studies to mitigate threats like retaliatory hunting and habitat loss. In Mexico, the Northern Jaguar Project offers ranchers up to US$1,200 monthly for photographic proof of jaguars on their land, incentivizing coexistence through financial rewards. In Belize, rural communities earn financial rewards for each new jaguar recorded, offering an alternative income source that values wildlife; payments can be 5 to 12 times higher than average daily earnings from other activities (Briggs-Gonzalez & Mazzotti, 2014).

In terms of more general policies related to habitat conversion Guatemala has certified forest concessions to prevent habitat loss while providing economic benefits, generating up to US$10 million annually from timber sales, with regions showing average jaguar densities of 11.28 per 100 km² (FSC, 2023; Polisar et al., 2017). The Jaguar Amazon REDD Project in Peru protects 180,000 hectares of jaguar habitat through carbon credits, supporting local communities, while the Jaguar Connection Project promotes self-sustainable conservation and community development using carbon credits across Latin America’s "Jaguar Corridor" (Hyde et al., 2022). In Brazil, the NGO Jaguar Conservation Fund (JCF) launched the “Jaguar Friendly Certificate” in 2000 to recognize ranches that protect jaguars, comply with environmental laws, and minimize wildlife conflict. Over 500,000 hectares are certified, enabling ranchers to access green financing and add value to meat and grain.

These successful examples show that ensuring the long-term survival of the species requires collaborative efforts between private and governmental sectors at federal, state, and municipal levels (De Marco et al., 2023). Brazilian law requires 20– 80% of native vegetation to be preserved on private lands, depending on the ecoregion (Brazil, 2012). Although, legal habitat protection exists in some ecoregions, the fragmentation and disturbed landscapes hinder jaguar dispersal and viable populations. Moreover — despite Brazil having a ‘national center for predators’ for one third of a century — lacks public policies to address jaguar–rancher conflicts.

While jaguars — and the whole Brazilian biodiversity — are owned by the state, Brazil is the single country within the G20 that has no policy of compensation such as PES for biodiversity (Ravenelle & Nyhus, 2017). This lack of clear policies leads to a dire scenario for the species that is functionally extinct in the Atlantic Forest and killed on sight throughout most of the remaining portions of their Brazilian range. For instance, in a single study site in the southern Amazon, at least 140 pumas and jaguars were killed in one year due to conflict with cattle ranchers (Michalski et al., 2006).

Paradoxically, for the last half-a-century Brazilian law (Brazil, 1998) prohibits wildlife management without specific plans authorized by the environmental agencies, which have never been issued on a large scale and typically are exceptional “case studies” focused on local perspectives (Marioni et al., 2013). Consequently, ranchers facing jaguar issues have no legal ways to control or manage the livestock predation. Once again, the lack of management for any wildlife species is a *sui generis* situation of Brazilian policy, with no parallels in the developed or developing world within the G20 (Ravenelle & Nyhus, 2017). Therefore, the development of financial compensation mechanisms to minimize this conflict throughout the species’ range is paramount, as previously discussed (e.g., Cavalcanti et al., 2010; Silveira et al., 2004). In this context, it is also crucial to recognize that the presence and maintenance of this predator on private lands entail high costs, simultaneously attesting the high biodiversity of extensive natural areas with good environmental quality. Jaguars play a crucial role in controlling prey species, maintaining ecological balance, and consequently ensuring the conservation of water, vegetation, soil, and other environmental benefits favorable to society (Burke et al., 2019; Thornton et al., 2016).

Here we provide for the first time a more operational approach for defining and establishing JCUs at regional scales, coupling the analyses of occurrence and home range patterns for Jaguars with fine-scale data of landholdings in Central Brazil. We use the state of Mato Grosso do Sul (MS) as a model to perform these analyses, given the abundance of studies on jaguar occurrence and its historical conflicts with cattle ranchers, and a well-defined land ownership situation. However, our approach can be easily adapted to specific contexts across the species’ range. With JCUs defined, we explore environmental compensation scenarios to support landowners, reduce conflicts, and promote a replicable conservation framework.

## 2. METHODS

### 2.1. Study Region

Mato Grosso do Sul (MS) is in the south of Brazil’s Central-West Region and covers an area of 35.7 million hectares (ha). The State is covered by the savannahs of the Cerrado ecoregion to 61%, while 25% of the territory is in the Pantanal floodplain, and 14% by the Atlantic Forest remnants (Silva et al., 2010). The state represented 1.6% of Brazil’s economy in 2021 and is the 15^th^ largest state in Brazil (SEMADESC, 2024). The primary economic activity is intensive agriculture, as well as livestock, carried out in 71,164 economically active ranches (SICAR, 2022). Most landholdings (83%) are larger than 1,000 hectares, while smallholdings (less than 50 hectares) represent only 4% of the area. Indigenous lands, meanwhile, occupy around 2.5% of MS, and the 33 protected areas in the state cover 0.92% of its total area.

### 2.2. Defining JCUs

We used jaguar occurrence records from camera-trapping and GPS-collar monitoring studies from JCF, as well as updated literature, and integrated with the previously described JCUs (Nijhawan, 2012) to determine the current distribution of the species in MS in period of 2010 to 2022. To establish the current occurrence area of jaguars, a 5 km radius buffer was created around each jaguar record. These identified areas were considered as "source populations" of jaguars in MS. To prevent the inclusion of many records of dispersing individuals along extensive sink-areas of their range, as jaguars are known to disperse up to 1,600 km (Jaguar Conservation Fund, unpublished data) and can be recorded in areas where there is no population at all (Sanderson et al., 2021), we filtered our data to include only sections with breeding evidence when females with cubs were recorded (see Sanderson et al., 2002).

We then overlayed the total extent of jaguars in the three “source populations” to the limits of 14,197 private areas based on the SICAR and Mapbiomas database (www.brasil.mapbiomas.org - Coleção 6, 2024). Thus, we defined a JCU as encompassing all the area delimited by the perimeter of all private properties within the three previously defined jaguar source populations. The importance of this new operational definition for JCUs is that it provides a better-defined spatial unit to which public policies can be more effectively applied to provide ecological and environmental compensation accounting for economic activities.

After this delimitation, the land-use patterns within JCUs, was obtained from Mapbiomas. With this data it is possible to define native vegetation and productive areas by comparing the proportion of remnants of natural vegetation with the expected values from environmental law (20% for properties located within Cerrado ecoregion and 35% for properties in the Pantanal ecoregion). Finally, it was possible to estimate the number of jaguar individuals expected in these areas by multiplying the total area of the buffers by the average density of the species in the Pantanal, Atlantic Forest and Cerrado ecoregions (Table 1).

**Table 1.**
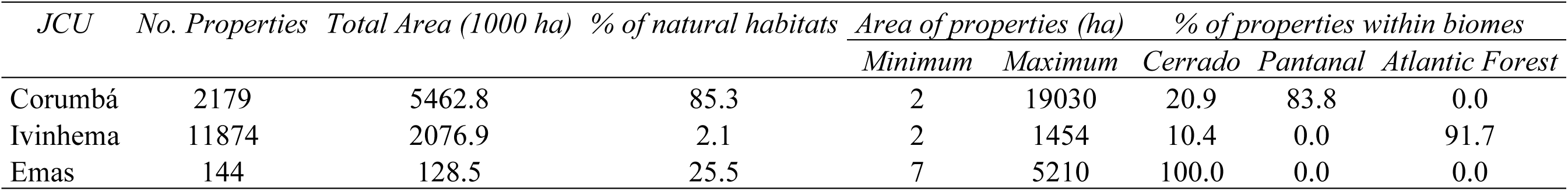
General characteristics of land use in the three JCUs defined for Mato Grosso State in Central Brazil, including number of rural properties, total area (in thousands of hectares- ha), % of natural habitats within JCUs, maximum and minimum area of rural properties and % of properties classified into the three biomes (Cerrado, Pantanal and Atlantic Forest) found in the State.

| *JCU* | *No. Properties* | *Total Area (1000 ha)* | *% of natural habitats* | *Area of properties (ha)* |  | *% of properties within biomes* |  |  |
| --- | --- | --- | --- | --- | --- | --- | --- | --- |
|  |  |  |  | *Minimum* | *Maximum* | *Cerrado* | *Pantanal* | *Atlantic Forest* |
| Corumbá | 2179 | 5462.8 | 85.3 | 2 | 19030 | 20.9 | 83.8 | 0.0 |
| Ivinhema | 11874 | 2076.9 | 2.1 | 2 | 1454 | 10.4 | 0.0 | 91.7 |
| Emas | 144 | 128.5 | 25.5 | 7 | 5210 | 100.0 | 0.0 | 0.0 |

### 2.3. Cost of jaguar predation on landholdings

One of the most important issues on jaguar loss refers to retaliatory killing of individuals by cattle ranchers (Cavalcanti et al., 2010). In the Pantanal, cattle make up 30–50% of the jaguar diet, mostly calves. Predation varies seasonally and decreases in wetter years due to greater wild prey availability (Tortato et al., 2015).

We built an Individual-Based Model (IBM) of predation based on parameters defined by Cavalcanti et al. (2010). In their study, it was established that predation of calves represents about 70% of livestock losses and that each jaguar kills a calf in intervals of 13.5 (± 15) days, and adults in intervals of about 25.5 (± 18.4) days. We defined the expected jaguar population within the JCUs by combining the total amount of area of natural remnants with estimates of density (Jędrzejewski et al., 2018), ranging from 0.9 to 4 individuals /100km² for different JCUs in the study area. For each jaguar in the JCUs, the IBM starts by deciding if a kill will be a calf or an adult (with the first with a 0.7 probability) and then sample a time interval for the next kill with the parameters given above. The process iterates over a year and is replicated 100 times for each individual jaguar to give the mean number of kills and the associated economic losses. The price of calves and adults killed at each time is also sampled from a normal distribution varying between ca. US$360.00 and US$450.00 for calves and ca. US$ 720.00 and US$830.00 for adults (see http://www.noticiasagricolas.com.br/cotacoes/boi). The entire process is summed across the jaguar population within the JCUs, giving the total amount of cattle ranch losses in a year and the economic loss due to predation.

## 3. RESULTS

### 3.1. Current distribution of jaguars in Mato Grosso do Sul and definition of JCUs

Our results allowed us to define three JCUs for MS (Figure 1), overlaying 43 municipalities of the State, with 21% of their areas encompassing a JCU, representing 54% of the total number of municipalities of the State. The three JCUs are defined based on the areas of a total of 14,197 rural properties and ranches with a total area of 7,671,000 ha, equivalent to 21.5% of the area of State (Table 1).

**Figure 1.**
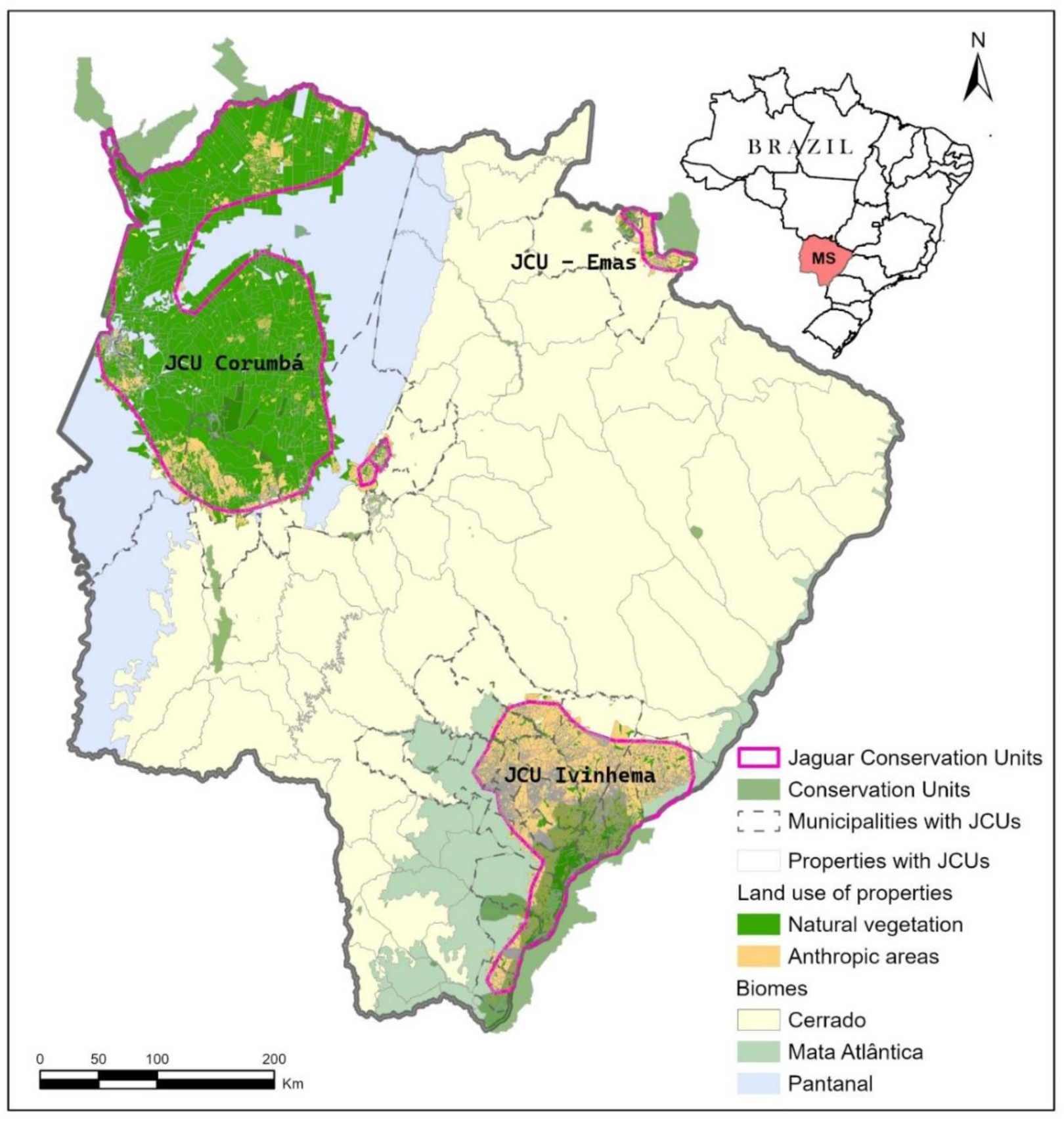
The three JCUs defined for Mato Grosso do Sul, based on data from Jaguar Conservation Fund (JCF) and JCUs database, expanded to encompass the limits of 14,197 rural properties in each region, from SICAR database.

Land-use analysis in JCUs reveals that natural vegetation of rural properties covers 5,145,500 ha (67% of the total area of the JCUs). However, the distribution of proportion of natural to total areas along the rural properties is highly skewed, with most properties with less than 10% of their areas preserved (Figure 2). It is therefore possible to evaluate if the relative amount of coverage by natural vegetation is in accordance with the Federal legislation. For those properties in the Pantanal ecoregion (mainly in Corumba JCU), there is a left-skewed distribution of natural cover, with most properties having more than 80% of natural cover within their area, and only 21% of their properties with less than the expected value of 35% of natural cover (Figure 2). The situation of the properties located in the Cerrado ecoregion (mainly Ivinhema and Emas JCUs), is entirely different, with a left skewed distribution in which only 37% of properties with more than 20% of natural cover.

**Figure 2.**
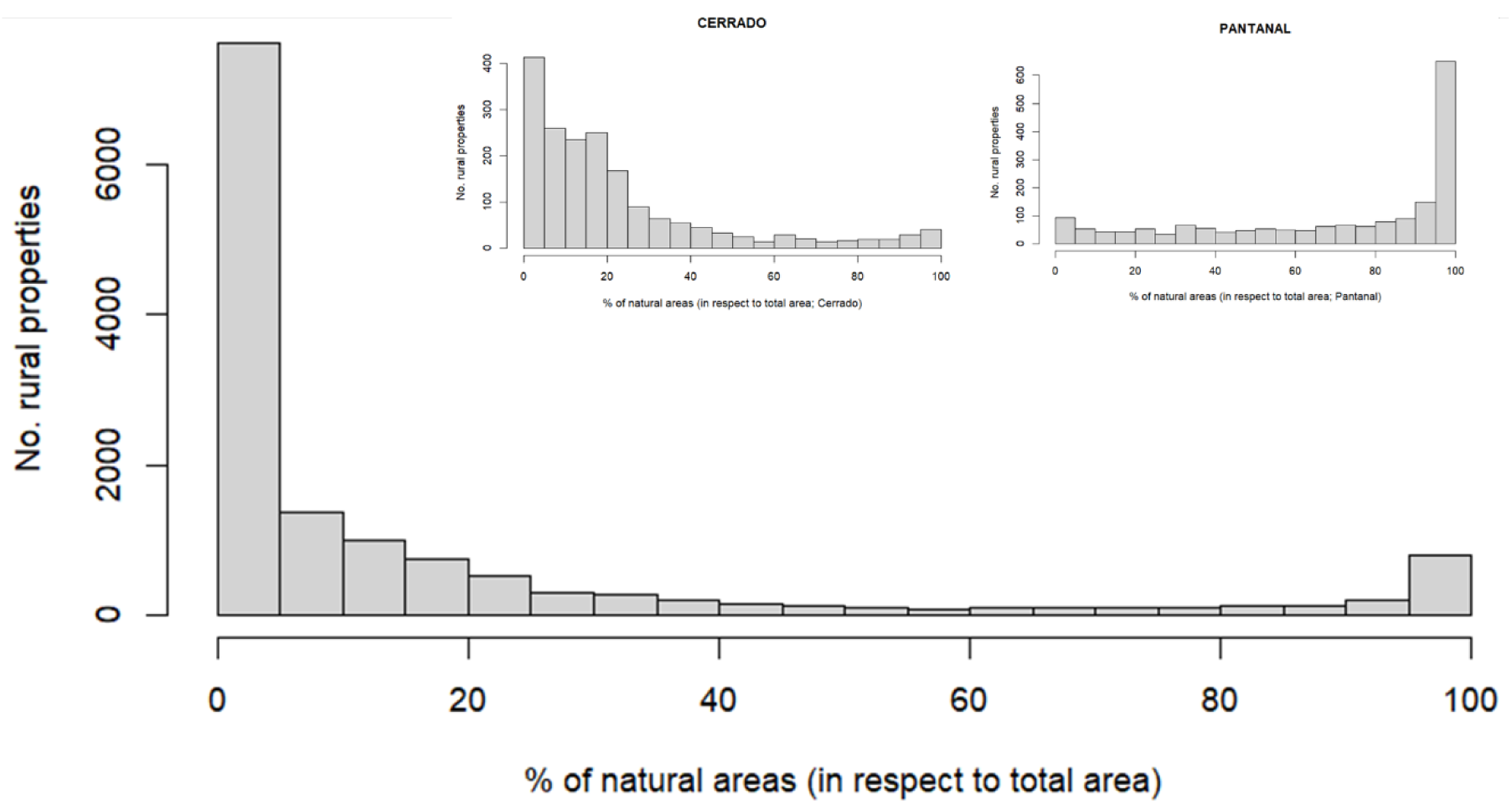
Distribution of proportion of natural habitats in rural properties for all JCUs in Mato Grosso do Sul, with inserts showing the same pattern for those properties in Pantanal biome (mainly Ivinhema JCU) and in Cerrado (Emas and Corumba).

These numbers also vary a lot among the three JCUs, as expected by the biome in which they are mainly located (Table 1). The Corumba JCU has a much larger proportion of natural habitats and larger rural properties, with only 24% of the rural properties not following the Forest Code. On the other hand, the landscape in Emas and Ivinhema JCUs, mainly found in the Cerrado and some remnants of Atlantic Forest biomes, is much more fragmented in terms of natural habitats and rural properties, with 55% and 87% of their properties with less natural cover than expected.

Thus, about 77% of properties in the JCUs have less natural cover than required by law, but these figures may be overestimated if some of the owners have extra- property areas to compensate for the local deficits. This figure depends on classification of natural versus anthropic areas at a more local scale, especially in the Cerrado. Even so, it is important to highlight that this deficit represents a smaller fraction (35%) in terms of total area combined for the three JCUs, as expected by considering the different levels of landscape fragmentation in the region and in the three JCUs. By considering only the 23% rural properties that are in accordance with the Federal legislation, their natural areas represent 96.4% of all natural areas in the region, in a total of 4,959,600 ha.

### 3.2. Kill patterns and Predation Costs

Our IBM starts by combining average estimates of population density for each JCUs, following Jędrzejewski et al. (2018) and assuming densities equal to 3 ± 0.5 individuals / 100 km² for rural properties in Corumba JCU, 1 ± 0.25 individuals / 100 km² for Ivinhema JCU, and 1.5 ± 0.5 individuals / 100 km² for Emas JCU. By combining values of density randomly sampled from these distributions with area of natural habitats in each JCU allows estimating the total population of jaguars in the region as about 1,457 individuals (CI 95% ranging from 983 to 1,813 individuals), and on average 96% of these would be found in Corumba JCUs.

Applying the IBM for predation on livestock based on this density distribution reveals that, on average, jaguar population within JCUs would kill about 24,800 individuals per year (CI 95% between ca. 17,400 and 32,500 individuals) (Figure 3A), with an average associated cost of US$13.1 million annually (CI 95% ranging from US$ 9.2 and US$ 17.2 millions) (Figure 3B).

**Figure 3.**
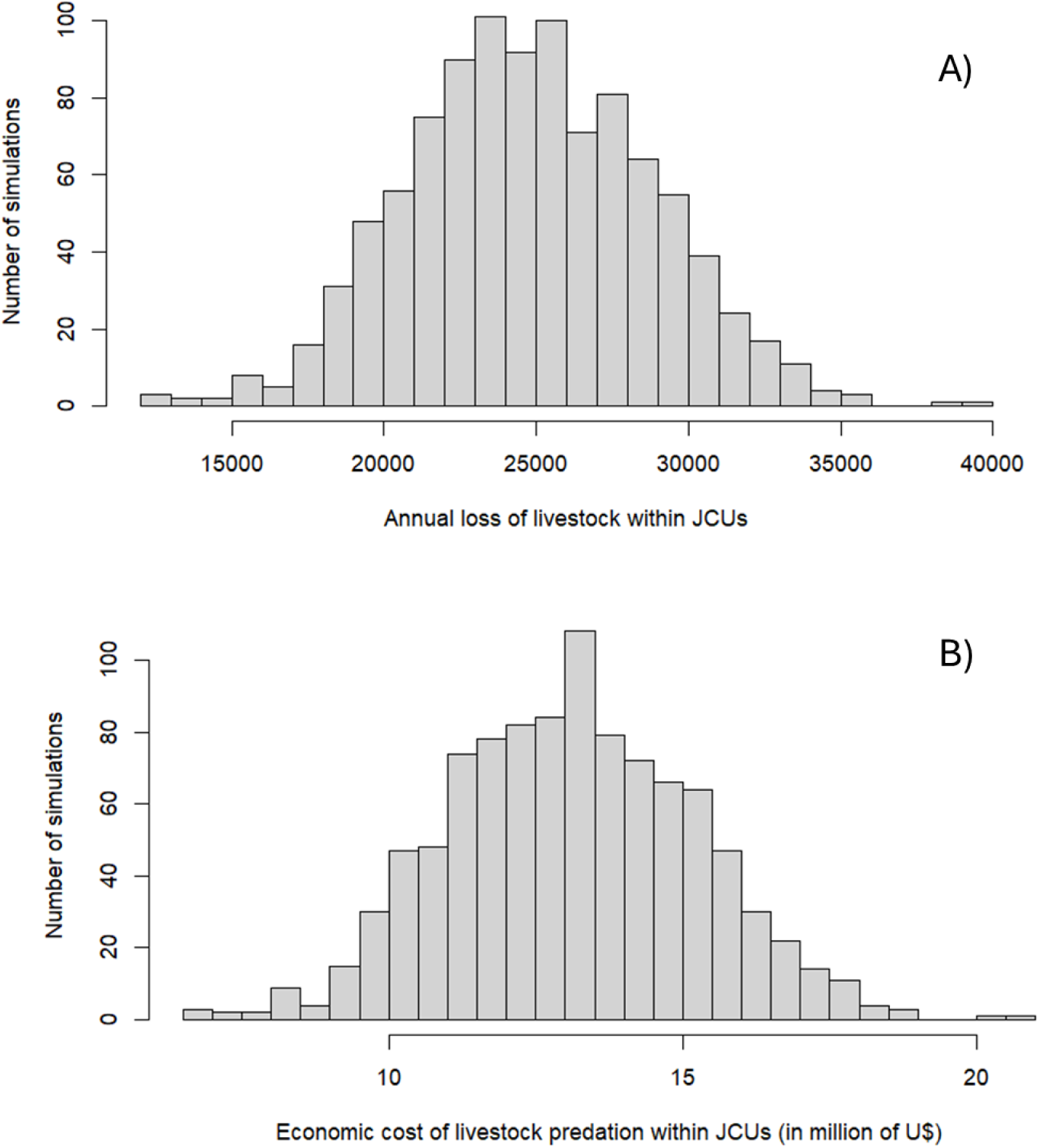
Statistical distributions obtained by the IBM simulating jaguar predation on livestock in the three JCUs, with the annual number of losses (A) and the economic costs of these losses, in US$ millions (B).

## 4. DISCUSSION

### 4.1. An operational definition of JCUs

In 2018, the United Nations Development Programme (UNDP) and partners launched the *Jaguar 2030 Roadmap* to strengthen jaguar conservation. The initiative incorporates JCUs as a foundation for conservation planning and prioritizes key landscapes, including the Pantanal (UNDP *et al*, 2018). Here we provide, for the first time, a more operational definition of JCUs, following the original reasoning by Sanderson et al. (2002). The JCUs are then redefined as the entire area of these rural properties overlapping with potential distribution of core jaguar populations. This new definition allows a more realistic discussion about the conservation strategies of populations in these regions, mainly involving PES and nature-based (economic) solutions.

In Mato Grosso do Sul’s Pantanal, extensive natural habitats support core jaguar populations amid cattle ranching on floodable grasslands, and most rural properties within the three JCUs (mostly concentrated in the Corumba JCU) have a relatively wide natural vegetation cover (Figure 1). That way, native vegetation used by core jaguar populations can still be reduced, highlighting the need for financial incentives to preserve current natural areas.

Despite legal vegetation requirements, properties within JCUs should be compensated for supporting jaguar populations. Landowners could be compensated based on the cost per hectare of maintaining a national park plus expected annual predation losses. According to a study conducted by the Brazilian Biodiversity Fund (Funbio, 2008), the annual maintenance costs of Brazilian national parks, excluding personnel expenses, totaled approximately US$8.51 million, resulting in an average of US$133,000 per park per year. Although these values were projected more than two decades ago, effective Park maintenance demands continuous investments in infrastructure, environmental monitoring, and the mitigation of threats such as wildfires and illegal anthropogenic activities. Updating these cost estimates is essential for the development of effective public policies and ensuring the long-term viability of biodiversity conservation in the Pantanal.

By considering only those rural properties that are in accordance with Forest Code, and using this reference value for the Pantanal region we could expect a total cost of US$5,120,000 for 320 thousand hectares of natural areas among the JCU’s. To this value should be added the expected annual cattle loss based on known regional predation rate. Therefore, the final value combines ranch hectares with expected herd loss in each ranch. It would be possible to select the benefited properties if stating that PES would apply only to properties that have no environmental passives, but because there are other economic incentives to follow the law, about 50% of the properties in our study site would be benefited by this policy.

Thus, using this reference value for the Pantanal region we could expect a total cost of US$83.8 million to support the JCUs. To this value should be added the expected annual cattle loss based on known regional predation rate, about US$13 million. It would be possible to select the benefited properties if stating that PES would apply only to properties that have no environmental passives, but because there are other economic incentives and advantages to follow the law, we expect that more landowners will be stimulated to regularize their legal situation. Notice that this mechanism of jaguar PES based on the total area of natural habitats within rural properties would be another incentive for regularization of the fiscal situation of rural properties and that, although a PES program would benefit only ca. 23% of landowners in the first instance, the total cost of the program would increase by about 30% even if all of them start to follow Federal legislation.

Despite appearing high, the cost supports conservation actions that benefit jaguars, other species, and habitats, offering ways to reconcile these costs with ranching and agriculture. The challenge lies in developing a systematic, permanent, and replicable mechanism on a scale suitable for Brazil’s reality. For instance, although losing 25,000 individuals by predation is indeed a large number, it represents only 0,1% of the total herd size estimated for the municipalities within the three JCUs (about 19 million heads), giving a first approximation of the overall magnitude of the cattle ranch economic activity in the region.

Moreover, in the context of PES, another advantage of economically reinforcing natural habitats within the JCUs is that it would ensure other environmental services and may at least in part compensate for the area’s high costs. Saeidl and Moraes (2000) estimated that the value of the ecosystem services and natural capital in Pantanal would achieve almost US$6,000.00 per ha/year, a value that could be even higher if more investments in other activities (e.g., ecotourism) is incentivized. Although this may seem “tangential” and indirect for the landowner, there is a worldwide trend to be more aware of these gains, which could be increased by carbon market regulation and other services in the future, thus largely compensating the cost of US$17.00 for maintaining the natural areas within JCUs. Also, some services have local impacts, while others provide broader environmental benefits to society.

The value estimated by Saeidl and Moraes (2000) is based on a general evaluation, but it would be important to more explicitly assess the potential direct and indirect environmental services provided by jaguars and the ecosystem functions performed by the species, as suggested by Burke et al. (2019) and Giozza (2018) developed a network of trophic interactions to identify the interrelationships present in the jaguar food chain, to arrive at a list of environmental services performed directly by the species, or indirectly by its prey or the food resources of its prey. Knowing the ecological functions performed by the jaguar, we sought scientific studies on the valuation of each ecosystem service provided by the species.

There are no records in the specialized scientific literature of studies valuing the jaguar for the environmental services it provides (Burke et al., 2019). For this reason, the economic value of the species by weighing up the costs and values associated with its presence in a given region, based on the valuation model proposed by Fredman (1995). According to this model, the total economic value of the species corresponds to its existence value summed with the value of its non-use (the right of future generations to know the species), finally added to its use value (environmental services). Jaguars contribute to three groups of ecosystem services (population control of prey species, regulation of vegetation structure, and cultural service), providing at least six ecosystem services. Through direct ecosystem services, jaguars can reduce damages to crops by regulating species such as peccaries. For instance, this regulation alleviates losses for farmers of US$1,921,008/year in the Emas National Park region (ICMBio/CENAP, 2016).

It’s also important to consider jaguars’ moral value, even if hard to quantify, as their protection is legally and ethically justified (Porfirio et al., 2014). In Brazil killing wildlife without permission can lead to a fine around US$900.00 per individual if the species is on official lists of endangered Brazilian wildlife, a very low value. This amount can be increased based on the circumstances of the offense. This is frequently the case of jaguars in Brazil in the few cases where the perpetrator is caught, and these fines amounted to a total of US$50,272 over a 12-year period, with the highest value imposed in 2022. Even so, they are relatively small when compared with the overall production gain and cattle losses.

### 4.2. Loss of Livestock by jaguar predation

Jaguar predation on livestock has long been recognized as a significant behavioral pattern in areas where they coexist with cattle. Balbuena-Serrano et al. (2021) estimated that approximately 18% of the jaguar current distribution in Brazil is at high risk of livestock depredation. Additionally, studies suggest that jaguars prey on approximately 0.6% to 2.3% of the total cattle herd on affected properties (Azevedo & Murray, 2007; Cavalcanti & Gese, 2010). Our IBM projects a loss due to predation within JCUs around US$13 million a year. Reimbursing such costs depends on the development of an economically viable model. Despite the challenges in implementing such a PES program, the value estimated by us is compelling enough to as a “shop window” demonstrate the magnitude of the loss of a species like the jaguar would produce, for the private sector to calculate the value of the PES. Properly recognize and proportionally share the concrete losses among the segments of society that contribute to the equation of jaguars’ existence is diffuse, belonging to many stakeholders and yet no one in particular (Brendin et al., 2015; Sandroni et al., 2022). This scenario usually leads to a “tragedy of the commons” scenario (Hardin et al., 2018).

Brazil’s law allows authorized hunting for conflict control (Law n°. 9,605/1998), but no permits for killing jaguars exist despite widespread illegal killings; few species are officially regulated as pests (Verdade & Campos, 2003; Fischer, 2018).

### 4.3. Economic opportunities for Jaguar PES programs

PES promotes human–predator coexistence by financially rewarding habitat conservation and jaguar presence. PES would financially incentivize tolerance for jaguars by focusing on the ecological benefits they bring, not just the economic losses. This could involve annual payments based on jaguar populations or rewards for documenting their presence. While challenges exist in implementing a large-scale PES program in Brazil, prioritizing areas with strong scientific data on jaguars and conflicts is a good first step. The flexibility of PES allows for adaptation to different regions, making it a viable long-term solution for jaguar conservation.

Every conservation initiative has an associated financial cost, which is why many implementation projects fail to foresee this cost in the long term. In the case of a jaguar PES program, actions aiming for coexistence can be targeted toward locations with persistent populations. This implies that the financial resources involved can be applied locally, not necessarily requiring significant amounts.

### 4.4. The Funding and Legislation Issues

Brazil has adopted the 2030 Agenda for Sustainable Development that includes the implementation of mechanisms for PES (IBGE, 2023). Similarly, the recent Global Biodiversity Framework for 2030 outlines the ensuring of Convention on Biological Diversity (CDB, 2022), a National Law 14,119 (Brasil, 2021) and Mato Grosso do Sul’s Law 5,235 (Mato Grosso do Sul, 2018) establish PES policies and programs for PES.

In 2021, the state government launched the first PES public notice, targeting rural landowners in the Formoso and Prata river basins, within the municipalities of Bonito and Jardim. The notice allocated US$171,427 to reward initiatives for forest conservation and restoration, as well as the conversion of degraded pastures and lands for more sustainable uses. The state government is the goal of expanding in the Serra da Bodoquena region and extending the program to other areas of the state.

Potential PES funding for jaguar conservation in MS includes private sources like the meat industry and ecotourism, and various government options. The meat industry is directly connected to the survival of jaguars in Brazil, as it plays a key role in driving both persecution of the species and, to a lesser extent, habitat loss. In this context, fees based on livestock losses to jaguars could fund local PES programs within JCUs. With Mato Grosso do Sul’s meatpacking industry processing 2.3 million cattle heads annually, a modest compensation fee of US$1.31 to US$1.63 per head would cover all predation losses within the JCUs. This approach would create a zero-sum scenario for ranchers, eliminating economic losses while allowing them to explore alternative uses for jaguar populations, such as ecotourism. Since this fee represents less than 1% of the current price per head (US$654-720), it offers a practical and economically viable PES solution to ensure the long-term survival of jaguars within the JCUs. In the end, this approach offsets losses and makes jaguars allies, positioning the meat industry as a biodiversity partner.

Ecotourism, on the other hand, has experienced a significant recent growth. In 2015, ecotourism in the Porto Jofre/MT region generated a minimum revenue of at least US$6,827,625/year (Tortato et al., 2017). Tourism focused on jaguar sightings has not been able to grow out of Pantanal, and, cannot be considered a source able to compensate for the whole of losses caused by jaguars in our JCUs, and even less so applicable to the whole of Brazil. Following Tortato et al. (2017), lobbying efforts failed to establish a jaguar PSA fund, limiting contributions from this sector.

The government can adapt various local initiatives to fund jaguar-focused PES programs in MS, drawing from multiple sources, like:

1. In the same fashion of the "Água DF" Program, a portion of state-level environmental service tariffs could be allocated specifically to jaguar conservation initiatives;
2. Adapting the Minas Gerais Environmental Services Policy framework (Minas Gerais, 2022) could provide a diverse funding pool, including public and private contributions, alongside traditional budget allocations;
3. Finally, exploring the Financial Market for Environmental Assets offers the potential to attract private investment through certified biodiversity instruments like jaguar conservation credits. These adaptable approaches can create a robust financial foundation for PES programs in MS.

Jaguar conservation in Brazil can be funded through national mechanisms like the National Environmental Fund, Fines Conversion Program, Diffuse Rights Fund, agribusiness and BNDES projects. Finally, the National Policy for PES, when regulated, could provide a framework for fundraising and various payment modalities to support jaguar conservation efforts.

### 4.5. Time-wise considerations

Institutional stakeholders broadly support jaguar conservation in central Brazil, but divergences exist regarding its implementation, on governance, hunting permission, and economic impact (Brendin *et al*., 2015). For this approach based on PSE to work in the long run, it is important to also provide guidelines for update and iterative improvements in the program. As the natural areas increase due to restoration targeting achieving legal parameters and owners are stimulated to avoid deforesting natural areas within their properties, it is expected that prey populations also increase and thus loss of livestock due to jaguar predation decreases.

On the other hand, it is essential to monitor these interactions, as jaguar populations and possible conflicts are expected to grow. JCUs should be periodically revised to add ranches and create new JCUs as protection and jaguar numbers increase. At the same time, revisions are needed as economic changes will alter land use, affecting compensation areas in JCUs and potentially impacting jaguar populations and demographics due to varying habitat use.

## 5. CONCLUDING REMARKS

To date, three legislative bills proposing financial compensation for losses caused by jaguars have been introduced in Brazil—one at the federal level and two at the state level. However, these proposals have not been enacted into law, likely due to broad scope, lack of technical criteria, and unclear payment standards. Here we propose a new operational definition of JCUs that could help protect core populations of jaguar by facilitating the application of PES and nature-based solutions.

From a more general perspective, the Kunming-Montreal Global Biodiversity Framework, adopted at Convention on Biological Diversity (CBD, 2022), sets an aim of living in harmony with nature by 2050, emphasizing actions to manage human-wildlife interactions effectively. While the concept of "coexistence" and “harmony” still lacks a clear functional and transcultural meaning, acknowledging the diverse impacts predators have on economic activities is crucial (IUCN, 2023; Pooley et al., 2021). This innovative defining JCUs paves the way for practical conservation strategies, fostering a balance between conservation and economic realities. By recognizing the role of jaguars in ecosystem services and aligning with global biodiversity goals, we move closer to "harmony" with nature - one that bridges cultural divisions and economic concerns with an eye toward sustainable coexistence.

## Acknowledgements

We would like to express our sincere gratitude to Caio Penido, President of IMAC – Instituto Matogrossense da Carne, and José Carlos Pedreira, member of COSAG – Conselho Superior do Agronegócio da FIESP, for their valuable insights and constructive feedback on the earlier drafts of this manuscript.

